# Semantic category and presentation frequency-based expectations are associated with distinct neural prediction effects

**DOI:** 10.64898/2026.05.11.724177

**Authors:** Margaret Jane Moore, Phuong Dang, XJ Ong, Jason B. Mattingley

**Author notes:** Corresponding Author: Margaret Jane Moore.

## Abstract

Past work has indicated that expectation can modulate neural responses to visual stimuli, but it is unclear whether these effects remain consistent across different types of unexpected stimuli. Here, we measured and compared neural prediction effects associated with semantic category and presentation frequency-based expectations in real-world object stimuli. Participants (n = 35) viewed real-world object images in rapid serial visual presentation (RSVP) streams. Semantically unexpected stimuli occurred when a stimulus was presented in a semantically incongruent stream. Low-frequency violations occurred when a rarely presented stimulus was displayed in a semantically congruent stream. Multivariate pattern analysis of electroencephalography (EEG) was used to quantify and compare the degree of information represented in neural activity for stimuli in different prediction conditions. Semantically expected stimuli yielded lower decoding accuracy relative to random (unpredictable) stimuli (125-313 ms post-onset) while semantically unexpected stimuli exhibited increased decoding accuracy (199-238 ms & 523-559 ms). Low-frequency violations yielded decoding accuracy which was not significantly different from semantically expected stimuli. Exploratory analyses indicated that dissimilarity between expected and presented stimuli quantified in terms of higher-level stimulus features, but not low-level visual features, modulated the observed neural prediction effects. These results demonstrate that different types of prediction violations have distinct modulatory effects on neural responses, providing novel insight into the neural implementation of predictive processing.

## Introduction

Predictive processing is theorized to play a fundamental role in enabling efficient perception of visual stimuli (Hogendoorn & Burkitt, 2019; Mumford, 1992; Press et al., 2020; Rao & Ballard, 1999; Summerfield & Egner, 2009). Predictive coding theories assert that incoming sensory information is compared with internal models of expected inputs and that this new information is differentially processed depending on whether it aligns with or violates expectations (Friston, 2005; Summerfield & Egner, 2009). Past work has provided evidence that predictive information modulates neural responses to visual stimuli in cases where statistical relationships provide information about the identity of upcoming stimulus features (Blom et al., 2020; Kok et al., 2014; Smout et al., 2019; Tang et al., 2018). However, it is not yet clear whether these effects remain consistent in cases where predictive relationships are defined in terms of broad semantic categories rather than learned associations between specific stimuli. Here, we used electroencephalography (EEG) to investigate how semantic expectations about upcoming stimulus identities modulate patterns of brain activity.

Past research has demonstrated that predictive information modulates how the brain responds to stimuli defined by elementary visual features (e.g. orientation, shape, location) (Ekman et al., 2017; Kok et al., 2014; Smout et al., 2019; Stokes et al., 2014; Tang et al., 2018). In these paradigms, expectation is manipulated by providing a cue or a sequence of previous stimuli which predict the identity of an upcoming stimulus, and brain responses are compared in cases where expectations are fulfilled or violated (Feuerriegel et al., 2021). While the occurrence and directionality of prediction effects have varied across different paradigms (den Ouden et al., 2023, 2024; Moore et al., 2024, 2025), a body of experimental work has reported reduced response amplitudes and reduced representational fidelity in expected relative to unexpected stimuli (Kaposvari et al., 2018; Richter et al., 2018; Smout et al., 2019; Summerfield & Egner, 2009). These results have been most consistent in paradigms where predictions were operationalised using stimuli with a single defining visual feature such as orientation or shape (Richter et al., 2018; Rideaux et al., 2025; Smout et al., 2019; Tang et al., 2018). By contrast, results have been less consistent in paradigms employing stimuli with both low-level visual and high-level semantic content, such as complex object or face images (den Ouden et al., 2023; Kaposvari et al., 2018; Moore et al., 2024, 2025). This variability implies that predictions may not be operationalised identically when the predicted stimuli engages both lower- and higher-level visual processing.

Most past predictive paradigms have generated statistical relationships in which the predicted stimulus was defined by a single feature (or set of features) which remained fixed across presentations. While this design is useful for simplifying predictive relationships for measurement in experimental contexts, it may not fully represent the complexity of predictive relationships that are calculated and employed by the brain in real-world environments (Ferrari et al., 2022; Hogendoorn & Burkitt, 2019; Press et al., 2020). In real-world contexts, the brain must generate complex expectations about broad categories of stimuli which are robust to noise due to differences in lower-level sensory input (e.g. variation in object orientation, lighting, occlusion, and precise identity) (Hohwy, 2020; Maarseveen et al., 2017; Moore et al., 2024). Past work has provided evidence that predictive processes influence perception of complex objects in real-world contexts (Spaak et al., 2022). However, it remains unclear whether the neural dynamics of predictive processing observed in simple prediction paradigms are consistent when the features associated with the predicted stimulus vary across presentations.

One approach for simplifying this issue to facilitate exploration in an experimental context is to generate predictions based on semantic categories. For example, if a person has an expectation that birds are commonly seen in trees, this prediction may be operationalised as expecting a specific semantic category (e.g. birds) rather than a specific set of lower-level features. Semantic categories do indeed tend to have some degree of commonality in low- and mid-level visual features, but variability in these attributes can be large across different exemplars (Carlson et al., 2013; Grootswagers, Robinson, & Carlson, 2019; Grootswagers et al., 2019). Moreover, past research has suggested that visual features cannot fully explain the patterns of neural activity that represent higher-level semantic concepts such as animacy (Grootswagers et al., 2019).

Considering semantic-level predictions also provides an interesting test of theories which imply that predictive processes should be largely analogous across multiple levels of the visual processing hierarchy (Rao & Ballard, 1999). If neural prediction effects are similar when predictions are operationalised in terms of lower-level features (e.g. orientation, shape) and higher-level features (e.g. semantic category), this would support this core implication (Kaposvari et al., 2018; Rao & Ballard, 1999). However, recent work has indicated that high-level and low-level feature violations may have differential modulatory impacts on neural responses to predictive relationships (Ferrari et al., 2022; Richter et al., 2018). Semantic-level predictive relationships offer a novel approach for exploring this issue, as stimuli that violate semantic expectations will inherently differ in their degree of dissimilarity from the expected semantic category. This dissimilarity can be quantified across a range of image features, ranging from high-level semantic distances to low- and mid-level visual features (Budanitsky & Hirst, 2001; T. Carlson et al., 2013; Grootswagers, Robinson, Shatek, et al., 2019; Victoria et al., 2019). If dissimilarity in terms of a specific feature-level strongly modulates observed neural prediction effects, this may provide insight into the specific level (or levels) of visual processing responsible for driving observed neural dynamics.

Semantic-level prediction paradigms also enable direct comparisons between expectations based on sematic category and expectations based on presentation frequency. For example, an object may be unexpected because it appears in an incongruent semantic context, or it can be unexpected because it is an uncommon exemplar of the expected semantic category. Attention effects related to presentation frequency or salience have been posited to have separable modulatory effects on neural responses relative to prediction violations, but this remains a contentious issue (Alink & Blank, 2021; Press et al., 2020; Rungratsameetaweemana & Serences, 2019). For example, past work has suggested that infrequent (i.e. unexpected) stimuli drive early attentional orienting responses which are distinct from predictive processes (Alink & Blank, 2021; Feuerriegel et al., 2021). However, it is not yet clear how this theorised early attentional orienting can be distinguished from neural responses indexing core predictive processes. Here we aimed to provide insight into the extent to which different types of prediction violation may differentially contribute to dynamic neural responses, and whether responses related to presentation frequency are distinct to those indexing expectation.

In the current study we evaluated how semantic category predictions modulate the representational fidelity of stimuli in cases where individual items either conform to, or violate, semantic expectations. We used multivariate decoding of neural activity patterns recorded with electroencephalography (EEG) to measure representations of familiar real-world objects in cases where they were semantically expected, unexpected, or random. Multivariate decoding is a data-driven approach which characterises complex activity patterns, capturing categorical relationships between brain responses and presented stimuli without relying on user-defined models (T. A. Carlson et al., 2020; Oosterhof et al., 2016; Robinson et al., 2023). This approach enables the measurement of stimulus-specific differences in information fidelity which may not be effectively captured by traditional univariate EEG analyses (Amado & Kovács, 2016; Kaposvari et al., 2018). Multivariate decoding has been used in previous work on how prediction effects modulate representations of visual object stimuli (den Ouden et al., 2023, 2024; Moore et al., 2025; Whyte et al., 2020)

Here, we recorded EEG brain activity as participants viewed sequences of images in familiar semantic categories (birds, mammals, plants, furniture) in semantically expected contexts (e.g. a bird amongst other birds) or in semantically unexpected contexts (e.g. a bird amongst furniture). We employed multivariate pattern analysis to quantify the extent to which neural representations of individual stimuli were modulated by whether they were expected, unexpected, or unpredictable (i.e. random). We observed robust semantic prediction effects in neural responses. However, the directionality of these effects was not consistent across stimuli that appeared in incongruent semantic contexts, and stimuli that were less-commonly presented members of the expected semantic category. Overall, the findings provide novel insight into how predictive processing is operationalised in the brain.

## Methods

### Participants

Thirty-five participants (23 female, 3 left-handed, average age = 23.7 years (SD = 6.5, range = 18-54) were recruited from The University of Queensland and were compensated for their time at a rate of $20 per hour. All participants reported normal or corrected-to-normal vision and provided informed, written consent. The study was approved by The University of Queensland Human Research Ethics Committee (HREA 2020/HE003101).

### Paradigm

The study aimed to characterise neural representations of object images that were expected, unexpected, or random in terms of their semantic context. We employed real-world object images (n = 64, visual angle = 3°) obtained from www.pngimg.com in rapid serial visual presentation (RSVP) sequences presented at fixation. These stimuli have been shown to yield reliable, above-chance decoding in similar EEG RSVP paradigms (Grootswagers et al., 2017; Grootswagers, Robinson, & Carlson, 2019). In total, 64 object stimuli were presented, but a subset of 20 stimuli (5 from each semantic category) were used to support decoding analyses. This approach was used to maximise per-stimulus trial counts in critical expectation conditions. Perceptual expectations were induced by manipulating the semantic category of objects presented in each stream. Specifically, semantically structured RSVP streams contained primarily objects belonging to one of four possible semantic categories: birds, mammals, furniture, and plants. Participants were instructed to view each of these RSVP sequences and count the number of red stars (n = 1-4) presented in each sequence as an orthogonal attentional probe task (Moore et al., 2024). All RSVP sequences used a 5Hz presentation rate (100ms exposure, 100ms ISI) and lasted approximately 36 seconds (180 stimulus exposures plus 1-4 attention probes).

Two different types of RSVP sequence were employed: Random and Semantically Structured. The purpose of the Random sequences was to generate the data needed to estimate neural representations for each object when no predictive semantic structure was present. In Random Sequences, all 20 stimuli included in decoding analyses were presented in random order (12 sequences, ∼120 exposures per stimulus).

The purpose of semantically structured sequences was to evaluate the extent to which neural representations are modulated by semantic expectations. Each semantically structured sequence was composed primarily of objects from a single semantic category (expected stimuli, approximately 90%), along with two different types of infrequent ‘unexpected’ stimuli: semantically unexpected, and low-frequency violations (60 streams, approximately 40 exposures per stimulus).

Semantically unexpected stimuli were objects that did not align with the present semantic category of the sequence within which they were embedded (e.g., a bird appearing in a plant sequence) (Figure 1, Panel C). These stimuli were selected from the 9 possible out-of-category stimuli (3 per semantic category) included in the decoding analyses. Each semantically structured sequence contained 16 semantically unexpected stimuli.

**Figure 1:**
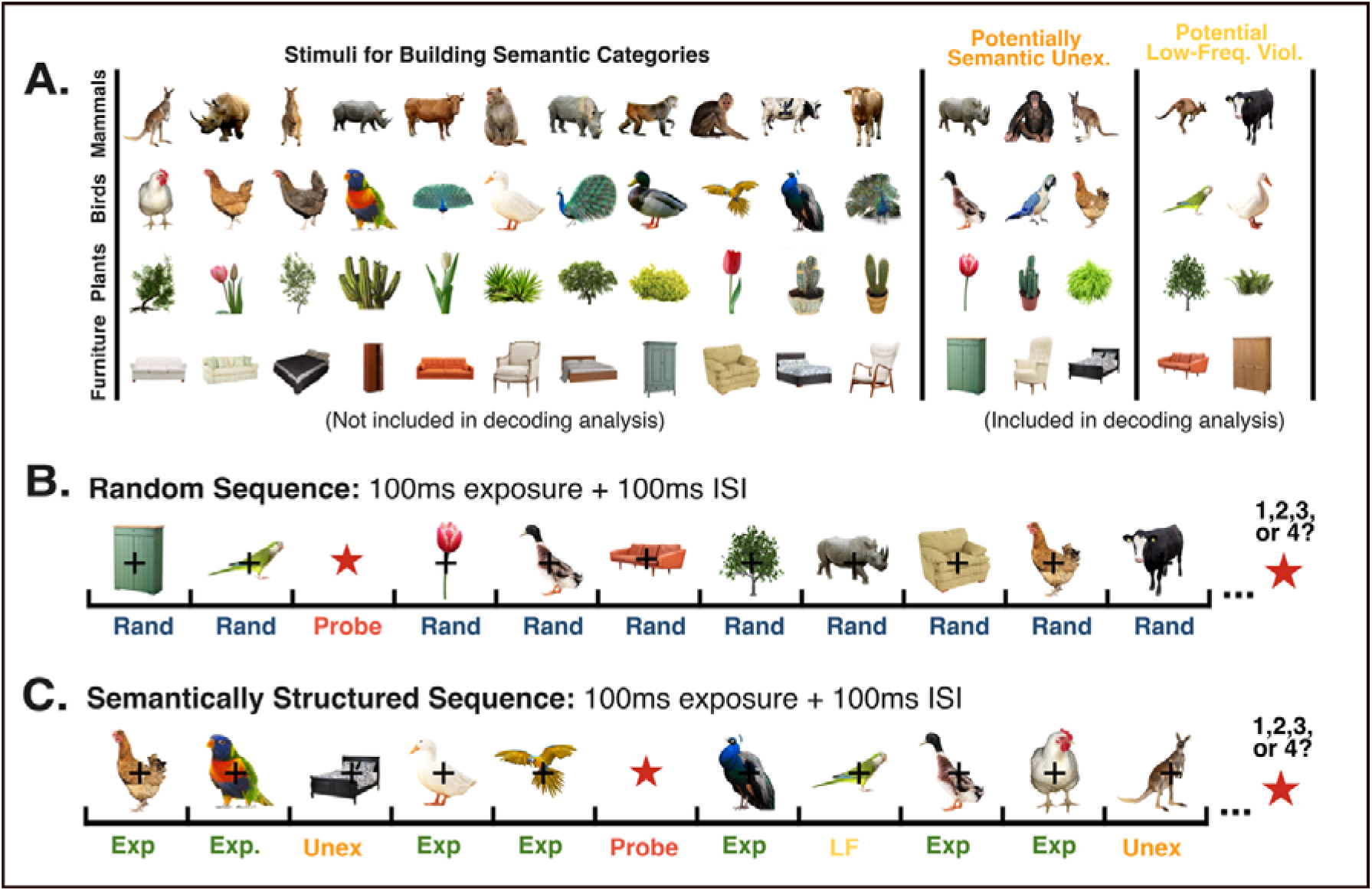
The stimulus set and experimental paradigm. A) All stimuli used in the experiment sorted into their respective semantic categories. Only a subset of the stimuli (labelled as potentially semantically unexpected or low-frequency violations) was included in decoding analyses. The assignment of stimuli to the semantically unexpected condition or low-frequency violations was randomised across participants. B) Example display of Random Block RSVP sequences. C) Example display of Semantically Structured Sequences. Semantically unexpected stimuli (Unex) occurred when a stimulus was presented in a semantically incongruent stream (e.g. bed appearing in a bird stream). Low-frequency (LF) violations occurred when an infrequent stimulus was displayed in a semantically congruent stream (e.g. rare green parrot appearing in bird stream). Each stimulus was on screen for 100ms (followed by a 100ms ISI). All object stimuli subtended 3° of visual angle. Participants were instructed to maintain fixation and to silently count the number of attention probes (red stars, 1-4) within each RSVP stream, and to report the number at the end of the trial.

Low-frequency violations were stimuli selected from the congruent semantic category but which were presented less frequently (Figure 1, Panel C). Two potential low-frequency violation stimuli were selected from each semantic category per participant and were presented a maximum of 4 times per RSVP sequence. This rate is lower than the presentation of all non-violation semantically expected stimuli that appeared in Semantically Structured sequences. Specifically, each of the 8 possible low-frequency violations was presented an average of 0.9 times per semantically structured block, while the remaining stimuli were presented an average of 4.0 times per semantically structured block. Low-frequency violations and semantically unexpected stimuli were pseudorandomly distributed within RSVP streams so that they were approximately evenly distributed, and so that these violations never occurred within in the first 20 stimuli in the stream.

Random and Semantically Structured RSVP streams were randomly intermixed in presentation order. The exact objects assigned to each possible violation category (e.g. low-frequency violations) was varied across participants.

Stimuli were centrally presented on a 120Hz monitor (1920 x 1080 pixels) at an approximate viewing distance of 60cm. Participants were instructed to maintain central fixation and provide responses using numbered keys on a standard computer keyboard. All experimental materials were coded in MATLAB using Psychtoolbox functions (Kleiner et al., 2007). The full paradigm lasted approximately 60 minutes, and participants were encouraged to take short rest breaks between sequences.

### EEG Procedure

Continuous EEG data were recorded using a BioSemi system and digitized at a sampling rate of 1024 Hz (64 electrodes, international 10-10 placement system). EEG data were pre-processed using EEGLAB (Delorme & Makeig, 2004). Specifically, raw EEG data were re-referenced to mastoid channels and were filtered using high pass (0.1 Hz) and low pass (100 Hz) frequency filters. Noisy electrode channels were identified using joint probability, and channels were rejected if they exceeded 5 standard deviations from the average (mean number interpolated = 1.23 electrodes, SD = 1.43, range = 0-5). These channels were reconstructed using spherical interpolation. EEG data were then down-sampled to 256 Hz, divided into stimulus presentation-locked epochs including the time interval from [-100 ms to +1000 ms] from stimulus presentation, and baseline corrected.

Central fixation was controlled and monitored using a video-based infra-red eye tracker (EyeLink 1000 Plus; SR Research). The eye tracker was calibrated at the beginning of the experiment using a standard nine-point calibration. RSVP streams only began after gaze was maintained for 500 ms on a central fixation cross presented at the beginning of each trial. If this criterion was not met within 3 seconds of fixation presentation, the calibration procedure was repeated. All EEG epochs in which an eye-blink occurred or in which fixation was not maintained within -200 to +200 ms relative to stimulus onset were excluded from subsequent decoding analyses. No other pre-processing or data cleaning was performed.

### Decoding Analyses

A multivariate pattern analysis (MVPA) regularised linear discriminant analysis (LDA) decoding pipeline was used to analyse the EEG data (Grootswagers et al., 2017; Oosterhof et al., 2016). This approach measures the discriminability of stimuli based on multivariate patterns of neural activity. Decoding analyses considered raw EEG voltages across all electrodes at each timepoint independently. Decoding analyses were performed in MATLAB using CoSMoMVPA functions (Oosterhof et al., 2016). All classifiers were trained and tested at the participant level, but overall results are reported at the group-level.

For each participant’s data, a pair-wise decoding classification scheme was applied across all unique stimulus pairs in the relevant condition of interest. Classifiers were trained to distinguish between each possible pair of stimuli using epochs where these objects were presented using a 5-fold cross validation scheme (trained on 80% of data, tested on 20% withheld data). Classifier performance was evaluated based on the average model accuracy within the withheld data across all cross-validation iterations. This process was repeated at each timepoint for each possible stimuli pair independently. Overall decoding accuracy is expressed as the average per-timepoint accuracy across all stimulus pairs and participants (chance accuracy = 50%). In this analysis, all classifiers were trained using data from Random sequences where no semantic expectations were present. These exact models were then applied to data from the key expectation conditions (random, semantically expected, semantically unexpected, low-frequency violations) to identify decoding accuracy differences across conditions. The number of trials included in each decoding model is reported in Supplementary Table 1.

### Statistical Inference

Statistical testing was conducted to evaluate whether decoding accuracy was above chance and to quantify differences in decoding accuracies across conditions. Wherever relevant, differences in decoding accuracy were evaluated by comparing the results to the decoding accuracy yielded by identical stimuli in random (i.e. unpredictable) sequences. This approach has been used in previous work to control for confounds arising from possible attentional, pupillary or domain-general differences between expected and unexpected stimuli (Feuerriegel et al., 2021; Moore et al., 2024, 2025). Bayes factors (BF) were calculated to represent the probability of the observed data occurring under the alternative hypothesis relative to the null hypothesis, for each timepoint independently (R package Bayes Factor) (Dienes, 2011; Rouder et al., 2009; Teichmann, 2022). All Bayesian analyses employed chance-level null hypotheses and alternative hypotheses with JZS prior (default scale factor = 0.707). In line with standard guidelines, Bayes Factors >10 were interpreted as strong evidence in support of the alternative hypothesis, and Bayes Factors <1/3 represented strong evidence in favour of the null hypothesis (Jarosz & Wiley, 2014; Rouder et al., 2009). Bayes Factors >3 and <10 were interpreted as representing moderate evidence in support of the alternative hypothesis.

To classify whether a specific time-window exhibited above-chance decoding, frequentist cluster-based permutation corrections for multiple comparisons are also reported. For these corrections, t-tests were conducted at each timepoint (one-tailed for vs. chance comparisons, two-tailed for between-model comparisons). Analyses yielding p-values <0.01 were included in clusters. Each comparison used 10,000 permutations to calculate the probability that each defined cluster (summarised by the cluster t-score sum) could occur when no underlying effect was present. All clusters with resultant p-values of <0.05 are reported.

Area under the curve analyses were used in cases where direct comparisons of time-point-level decoding accuracy was not feasible (e.g. exploratory analyses). Specifically, these analyses were used to compare the magnitude of observed neural prediction effects across different stimuli (where group-level sample sizes were low) and to compare decoding performance between different violation types where stimuli were not identical. For each comparison, area under the decoding curve was calculated for each participant (trapezoid integration method, 80-600ms) and compared with that of random stimuli (positive values = improved decoding vs. identical random stimuli, negative values = lower decoding vs. identical random stimuli). These area under the curve difference scores were used to compare decoding magnitude differences at a group level using Bayesian one-sample t-tests (versus zero) and two-sample t-tests (versus other expectation condition). Where relevant, these analyses were applied using a sliding-window approach to explore whether results were consistent at different timepoints following stimulus onset.

### Data Availability

All code and experimental materials associated with this study are openly available on the Open Science Framework (https://osf.io/s6k7u/). Due to size restrictions, EEG data is available upon request from the authors.

## Results

### Behavioural Results

All participants demonstrated above-chance (>25% correct) performance on the attention probe task (mean accuracy = 88.5%, SD = 10.6%, range = 52%-100%). Participants were able to maintain fixation with an average of 86.4% of trials meeting fixation inclusion criteria per-participant (range = 51.1% - 98.9%). All participants reported noticing the presence of the semantic structure and occasional violations of this structure during debriefing questions. However, no participants reported noticing differences in the frequency of presentation across different stimuli.

### Semantically expected stimuli yield reduced decoding accuracy

In our initial analysis of the EEG data we compared decoding accuracy for random, semantically expected, and semantically unexpected stimuli. Classifiers were trained to distinguish stimuli using data from Random sequence epochs. This classifier was then applied to data from expected and semantic violation stimuli appearing In Semantically Structured sequences. Decoding accuracy for random, semantically related (expected), and semantic violation stimuli was significantly above chance within the typically reported window for neural representation of visual objects in RSVP streams (80ms - 600ms, Carlson et al., 2013) (Supplementary Table 1) (Figure 2). Critically, decoding accuracy for semantically expected stimuli was lower than decoding accuracy for random stimuli between 125-313 ms (mean BF = 1.54 x 10^16^), whereas decoding accuracy for semantically unexpected stimuli was significantly higher than for random stimuli in time windows between 199-238 ms (mean BF = 2.07 x 10^3^) and between 523 – 559 ms (mean BF = 1.10 x 10^5^).

**Figure 2:**
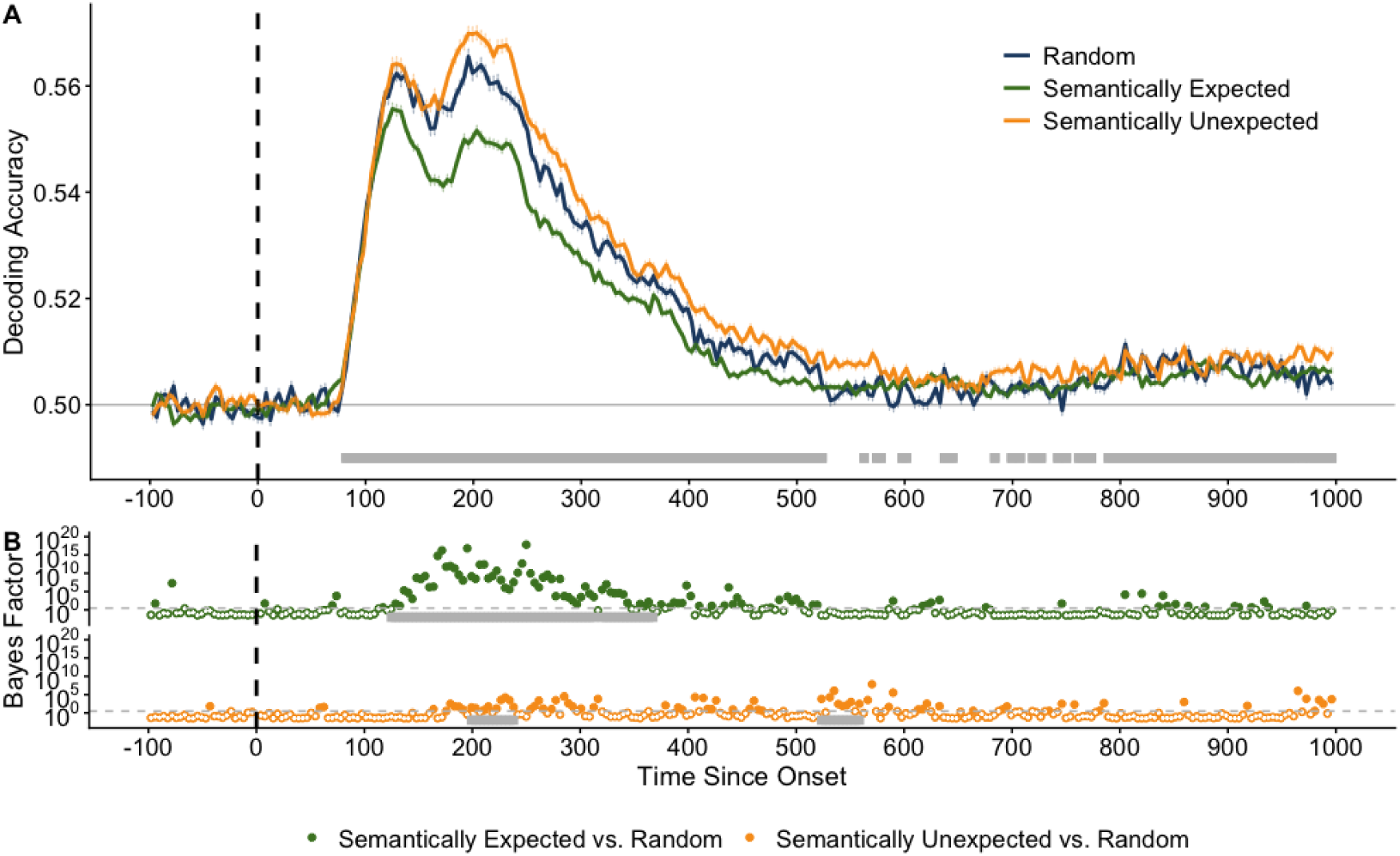
Decoding accuracy differences between semantically expected, semantically unexpected, and random stimuli. Grey bars denote regions where all visualised decoding curves are significantly above chance, as quantified by cluster-based permutation corrections for multiple comparisons. (A) Mean decoding accuracy for all participants across time (x-axis, in milliseconds). Stimulus onset (0ms) is denoted as a vertical dotted line and chance performance (50%) is represented in grey. (B) Bayes factor plots for between-model comparisons. The boundary for moderate evidence (BF = 3) is denoted by a horizontal dashed line. Tests yielding BF > 3 are represented by filled dots, while tests yielding BF < 3 are not filled. Time windows in which differences remained significant following cluster-based permutation corrections are highlighted in grey. All reported time points are rounded to the nearest millisecond.

### Low-frequency violations and semantically expected stimuli have comparable decoding accuracy

Next, decoding accuracy for random and low-frequency violation stimuli were compared. Recall that these low-frequency stimuli always matched the semantic category of the stream within which they appeared. Classifiers were trained to distinguish between low-frequency violation stimuli using data from the random sequences. This classifier was then tested on data from low-frequency stimuli occurring within semantically structured sequences. Notably, decoding performance was not compared across different semantic expectation/violation conditions in this analysis as stimuli used for low-frequency violations were always semantically related to the sequences within which they appeared. Decoding accuracy for random and low-frequency violation stimuli was significantly above chance within the expected window for neural representation of visual objects in RSVP streams (80ms - 600ms, Carlson et al., 2013). Decoding accuracy for low-frequency violations was significantly lower than accuracy for random stimuli between 74-516 ms (mean BF = 3.95x10^7^) post-onset (Figure 3).

**Figure 3:**
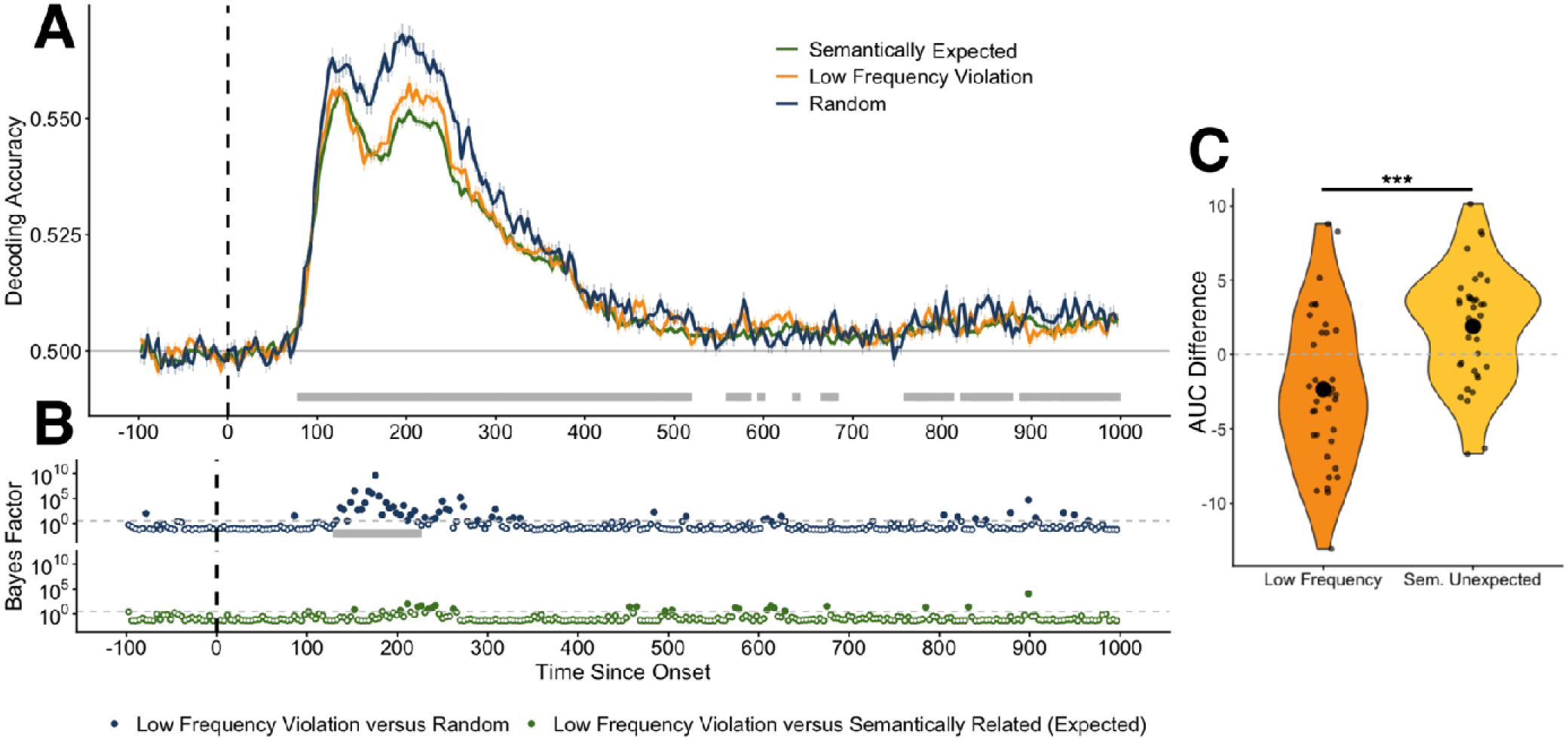
Decoding accuracy differences between semantically related (expected), semantically unexpected, and random stimuli. Grey bars denote regions where all visualised decoding curves are significantly above chance, as quantified by cluster-based permutation corrections for multiple comparisons. (A) Mean decoding accuracy for all participants across time (x-axis, in milliseconds). Stimulus onset (0ms) is denoted as a vertical dotted line and chance performance (50%) is represented in grey. (B) Bayes factor plots for between-model comparisons. The boundary for moderate evidence (BF = 3) is denoted by a horizontal dashed line. Tests yielding BF > 3 are represented by filled dots, while tests yielding BF < 3 are not filled. Time windows in which differences remained significant following cluster-based permutation corrections are highlighted in grey. All reported time points are rounded to the nearest millisecond. (C) Area between the curves analysis results. AUC Difference corresponds to the difference in area under the curve for low frequency violations (versus decoding accuracy in random sequences) and semantically unexpected stimuli (versus decoding accuracy in random sequences, as visualised in Figure 2). Individual-level values are reported as small dots and condition means are depicted as large points. Violin plot areas summarise the AUC difference distributions in each condition. *** = BF > 30.

**Figure 4:**
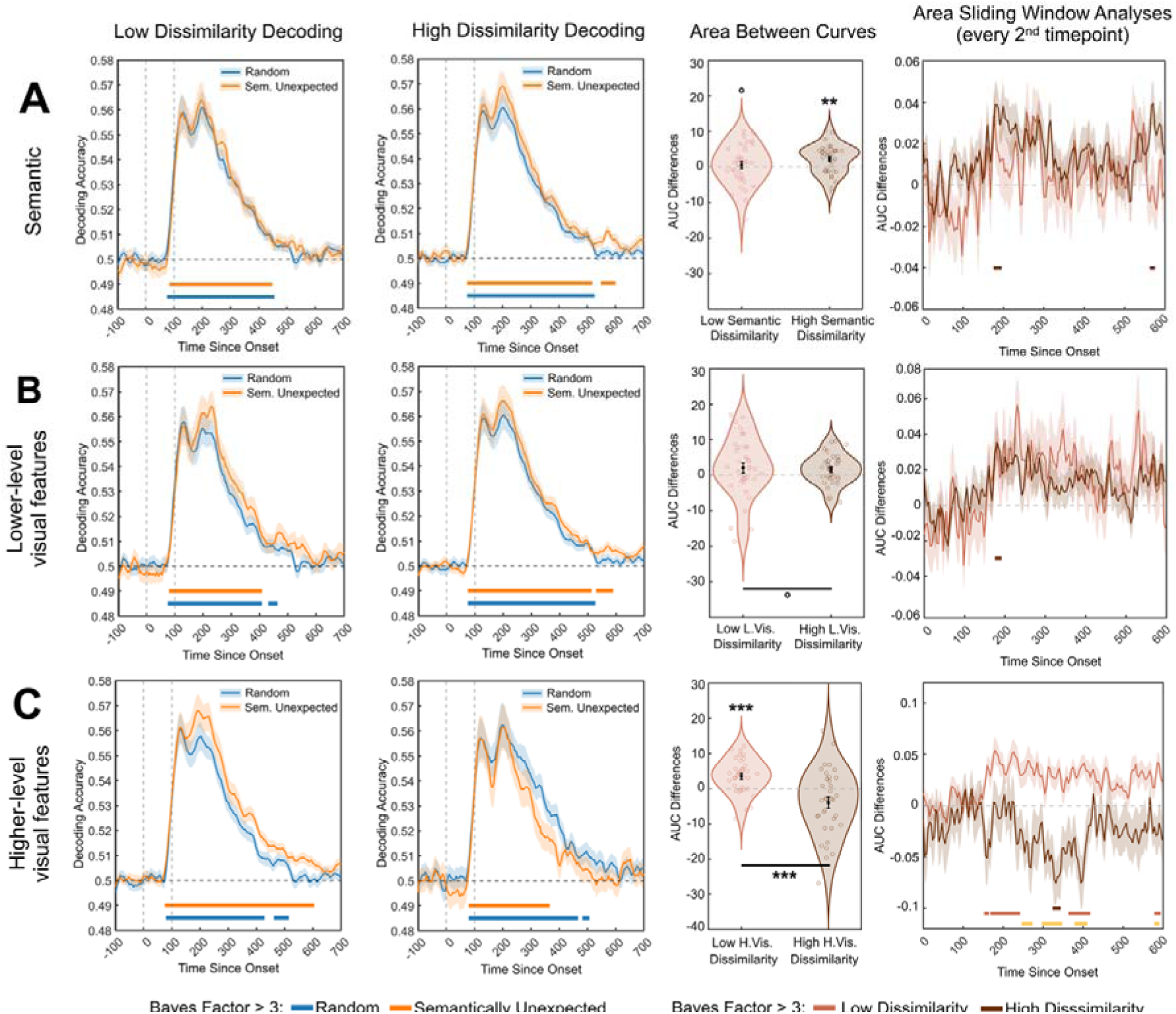
A comparison of decoding results within stimuli with high versus low dissimilarity relative to the sequences they were presented in. Results for comparisons based on (A) semantic distance, (B) low-level visual feature dissimilarity, and (C) high-level visual feature dissimilarity are visualised. Each row presents decoding performance in stimuli with low versus high dissimilarity scores and area between the curves analysis results. In decoding plots, mean decoding accuracy across time (x-axis, in milliseconds). Stimulus onset (0ms) and offset (100ms) are denoted as vertical dotted lines. Chance performance (50%) is marked by horizontal grey lines. Coloured bars denote time windows where decoding performance is above chance (BF > 3). In AUC comparison plots, violin plots visualise the distribution of participant-level differences in the area between decoding curves across dissimilarity conditions between 0-600ms after onset. Dissimilarity group is reported on the x-axis, and the y-axis reports the area under the curve for semantically unexpected stimuli minus the area under the curve for random stimuli. Dissimilarity group mean is represented as a dot with standard error of the mean represented by the error bars. Results for Bayesian one-sample t-tests comparing each group distribution to zero are reported in or above each relevant distribution, while results for Bayesian t-tests comparing across dissimilarity conditions are denoted between group distributions. Area Time Series plots report sliding time-window analyses (mean of 2 neighbouring timepoints) reporting AUC differences in unexpected versus random decoding accuracy across dissimilarity groups. Coloured bars denote time windows where one-sample t-tests (versus zero) yield BF >3. BF >3. ◦ *= BF <1/3, ** = BF > 10, *** = BF > 30, unspecified = inconclusive*.

Decoding accuracy for low-frequency violations was also compared with that for semantically expected stimuli. This comparison should be interpreted with caution as the stimuli in each condition were not identical within individuals (see Figure 1). However, there were no statistically significant differences in decoding accuracy between low frequency violations and semantically expected stimuli. Across the entire time window considered, decoding differences between these two conditions had a maximum of two consecutive timepoints yielding BFs > 10 (commencing at 223, 238, and 613 ms post-onset).

Area-between-the-curve analyses were also conducted to compare decoding performance differences between non-identical stimuli. One-sample Bayesian t-tests provided moderate evidence in support of higher decoding accuracy for semantically unexpected versus random stimuli (Mean AUC difference = 1.90, BF = 5.74), as well as moderate evidence of lower decoding accuracy for low-frequency violations relative to random (Mean AUC difference = -2.35, BF = 3.81). However, there was also strong evidence for a difference in decoding (relative to random) between low-frequency violation and semantically unexpected stimuli (BF = 99.5, Figure 3, Panel C).

### Lower- and higher-level image features differentially contribute to prediction dynamics

Next, exploratory analyses were conducted to provide preliminary insight into the extent to which the representational dynamics of predictive processing effects may be driven by different levels of stimulus features. This analysis was conducted because semantically unexpected stimuli differed from the semantic category they were presented in, both in terms of visual features and semantic labels. These analyses explored the relative contribution of these different information sources to the observed elevated decoding accuracy in semantically unexpected stimuli relative to random stimuli. The analysis was conducted using dissimilarity metrics assigned through semantic label and visual feature dissimilarity measures. Across all metrics, the average dissimilarity between individual stimuli and the semantic category they appeared in was represented by the average dissimilarity between each exemplar and each member of the relevant semantic category.

Semantic label dissimilarity was quantified by measuring the semantic distance (1 – cosine similarity) between each stimulus name (e.g. duck, closet, cactus) using a continuous Skip-gram model implemented in MALAB (Mikolov et al., 2013). Visual dissimilarity was quantified using neural network layer extractions generated by the deep convolutional neural network VGG-16 (Simonyan & Zisserman, 2014). Dissimilarity was measured using the Euclidean distance between VGG-16 layer activations for each pair of stimuli. This measurement was repeated at two different levels of VGG-16 activation, a lower-level visual layer (conv1_1) corresponding to orientation and edge features and a higher-level visual layer (fc7) corresponding to abstract, high-level image descriptors, prior to identify classification. These dissimilarity measures were adopted to explore how different hierarchical levels of processing may contribute to the observed neural prediction dynamics.

Median splits were employed to group each stimulus as low dissimilarity or high dissimilarity based on each considered dissimilarity metric. Decoding analyses comparing classifier performance for semantically random and unexpected stimuli were repeated for each stimulus subgroup. If the magnitude of neural prediction effects is not different across dissimilarity comparison groups, this would suggest that different processing levels may contribute equally to the observed neural decoding effects. By contrast, if significant differences are observed across dissimilarity comparison groups, this would provide some insight into which specific stimulus features might be driving neural responses.

The dissimilarity measures exhibited weak but significant correlations. Semantic dissimilarity was weakly correlated with both low-level (r = 0.24, R^2^ = 0.06, p < 0.001) and high-level visual feature dissimilarity (r = -0.14, R^2^ = 0.02, p < 0.001), and dissimilarity scores were weakly negatively correlated between low- and high-level visual features (r = 0.18, R^2^ = 0.03, p < 0.001).

Differential decoding accuracy patterns emerged when stimuli were grouped according to different dissimilarity metrics. When stimuli were grouped according to semantic label dissimilarity, there was strong evidence for decoding differences between unexpected and random stimuli within stimuli with high dissimilarity scores (*BF* = 10.51, mu ≠ 0), but strong evidence *against* decoding differences in stimuli with low-dissimilarity scores (*BF*= 0.21, mu ≠ 0). Direct comparisons of difference magnitude across these dissimilarity groups yielded inconclusive evidence (BF = 1.04). Sliding window analyses indicated that effects at the level of the full considered time epoch were likely driven by intermittent increases in decoding differences within high dissimilarity stimuli (relative to zero) between 179–191 ms (4 timepoints) and 566–570 ms (2 timepoints) post-onset.

When stimuli were grouped according to low-level visual feature dissimilarity, no significant differences in decoding accuracy emerged in low-dissimilarity (*BF* = 0.46, mu ≠ 0), high-dissimilarity (one-sample *BF* = 1.70, mu ≠ 0), or between dissimilarity groups (BF = 0.19). However, sliding window analyses identified intermittent evidence for differences between unexpected and random decoding performance in only high dissimilarity stimuli at 179–187 ms (3 timepoints) post-onset.

When stimuli were grouped according to high-level visual feature dissimilarity, there was strong evidence for improved decoding accuracy for semantically unexpected stimuli (relative to random) within low dissimilarity stimuli (*BF* = 201.29, mu ≠ 0), but inconclusive evidence in support of a difference from zero in high dissimilarity stimuli (*BF* = 2.28, mu ≠ 0). This comparison yielded strong evidence of a difference in the relative magnitude of decoding performances between unexpected and random stimuli in high- and low-dissimilarity groups (BF = 329.81). Sliding time-window analyses indicated that this difference emerged at 250 ms and continued intermittently through subsequent time points in the considered decoding window.

## Discussion

This study aimed to explore how semantic category predictions modulate the representational fidelity of stimuli in cases where individual items either conform to, or violate, semantic expectations. The results of this project provide novel insight into the neural dynamics associated with semantic-level predictive relationships. Participants were able to reliably complete the behavioural task and semantically expected stimuli yielded lower decoding accuracy relative to random stimuli throughout the time course generally associated with visual object representation (Carlson et al., 2013). Conversely, semantically unexpected stimuli exhibited increased decoding accuracy. Low-frequency violation stimuli were used to explore whether neural dynamics were consistent across different types of prediction violations, and these stimuli yielded decoding accuracy which was not significantly different from that of semantically expected stimuli. Additionally, exploratory analyses indicated that dissimilarity in terms of higher-level stimulus features, rather than low-level visual features, modulated the observed neural prediction effects. Taken together, these results provide novel insight into the neural implementation of predictive processing while suggesting several key considerations for future research,

### Evidence for expectation-suppression of neural responses in complex object stimuli

Decoding accuracy for semantically expected stimuli was comparatively lower than decoding accuracy for random stimuli. This finding aligns with “expectation suppression”, the pattern of neural responses that predictive coding theory posits to represent attenuated responses to incoming sensory information that aligns with internal models of expected stimuli. Predictive coding is a highly influential theory detailing how predictive processing might be implemented at a neural level (Rao & Ballard, 1999). However, evidence supporting expectation suppression as a core feature of predictive processing (and evidence for predictive coding accounts over competing explanations) is not entirely consistent (Feuerriegel et al., 2021). Notably, studies using complex object stimuli (like those used here) have typically found no evidence for neural prediction effects, or prediction effects that are not in line with expectation suppression (den Ouden et al., 2023; Moore et al., 2024, 2025). In the context of the present study, this indicates that the specific way that predictive relationships are operationalised may be an important determinant of associated neural responses. Neural prediction effects which are not in line with expectation suppression-like effects have been identified when similar stimuli were used in probabilistic cueing designs (Moore et al., 2024, 2025), but a different pattern emerged in the current study. There are several paradigm-specific factors which may help account for these differences.

The attentional salience of prediction violations might have been higher in our study relative to past work using similar stimuli. For example, semantically unexpected stimuli in our study involved comparatively salient mismatches between individual stimuli and their semantic context, while previous studies used probabilistic cueing violations which participants were not consciously aware of (Moore et al., 2024, 2025). It is possible that violations (or predictive structures) that elicit conscious awareness generate more pronounced (or qualitatively different) neural responses due to increased top-down processing and conscious surprise (Alink & Blank, 2021; Feuerriegel et al., 2021). Past research has also indicated that task demands may modulate the occurrence of prediction effects (Auksztulewicz et al., 2017; Smout et al., 2019; Stokes et al., 2014). Participants’ degree of exposure to predictive relationships has also been identified as a key factor impacting the occurrence of neural prediction effects (Ferrari et al., 2022). Notably, recent work has indicated that predictive relationships may modulate perceptual decision-making performance instead of sensory encoding processes (Gastrell et al., 2026.) In the present study, no responses to predictable stimuli were provided, and observed neural effects emerged in time windows associated with early visual processing stages, making it unlikely that the effects observed here were confined to perceptual decision-making processes. The current study is in line with past work indicating that the occurrence and directionality of neural prediction effects are dependent on paradigm-specific features. Future work is needed to more explicitly identify specific factors that modulate the occurrence/directionality of prediction effects.

### Different violation types are associated with different neural response patterns

Notably, low frequency stimuli yielded comparable decoding accuracy to semantically expected stimuli. This statistical result should be interpreted with caution as the exact stimuli compared in these conditions were identical at the group level, but not within individual participants. However, even in the absence of direct statistical comparisons, decoding accuracy for low-frequency stimuli showed a clear, qualitative difference relative to decoding accuracy for semantically unexpected stimuli. This result is theoretically important because it suggests that not all types of prediction violation have analogous modulatory effects on neural activity. This conclusion is in line with previous literature documenting variation in the occurrence and directionality of neural prediction effects across different paradigms and methods for operationalising predictive relationships (den Ouden et al., 2023; Hogendoorn & Burkitt, 2018; Moore et al., 2025; Tang et al., 2018). The result also suggests that, in cases where a stimulus’ broad semantic category conforms to expectations, variability in the specific combinations of features which define the item’s unique identity may not be detected as prediction violations. This approach to operationalising predictions based on high-level categories may help enable efficient perception in real-world scenarios where expected sensory input may be noisy or not entirely visible (Spaak et al., 2022).

The possibility that predictions may be operationalised based on semantic categories (rather than stimulus-specific information) has implications for future paradigms aiming to tap predictive processes. For example, in a probabilistic cueing design where a cue indicates that the subsequent stimulus is likely to be from a specific semantic category, it is possible that ‘unexpected’ exposures of other stimuli from within the same semantic category may not activate comparable predictive neural responses compared with unexpected stimuli from outside the relevant category. In this context, future studies should consider the extent to which the degree of similarity between expected and unexpected stimuli may be responsible for driving the occurrence and magnitude of neural prediction effects in complex stimuli.

One possible explanation of why the neural response pattern observed for low frequency stimuli was not aligned with that observed for semantical violation stimuli is that the frequency manipulation used here was not strong enough to cause low-frequency stimuli to be encoded as being uncommon. Here, each possible low-frequency stimulus was presented (on average) less than one time per semantically structured block, or roughly 4.5 times less frequently than the other analysed stimuli. It is plausible that even rarer stimulus presentations (e.g., once per experiment) may be required to elicit robust low-frequency violation effects, but such a design would not be compatible with the multivariate pattern analyses used here. It is also plausible that a greater degree of attention is needed to detect low-frequency violations relative to semantically unexpected stimuli. Past literature has indicated that the degree of task-related attention required can modulate the occurrence of neural prediction effects (Auksztulewicz et al., 2017; Smout et al., 2019). The attentional-probe behavioural task we employed here did not necessarily tax attention to stimulus features, which could plausibly have prevented low-frequency stimuli from being attended to the extent necessary to facilitate detection of their rareness. Overall, the results presented here suggest that different types of prediction violations may elicit differential effects on neural responses. Future research is needed to test whether this result remains when lower-frequency violations that are more task-relevant are considered.

### Prediction effects may be driven by specific stimulus features

Our exploratory analyses of the stimulus factors driving neural prediction effects provided preliminary evidence that neural prediction effects may not be equally modulated by all presented stimulus features. When stimuli were semantically unexpected, the magnitude of prediction effects was modulated by dissimilarity in terms of higher-level visual features, but not lower-level visual features. Decoding differences were also present in stimuli with high-, but not low-semantic label dissimilarity scores. Notably, this finding aligns with previous fMRI work indicating that prediction errors may be modulated by violations in expected high-level features, but not low-level features (Richter et al., 2024). However, it is worth noting that this previous paradigm required participants to perform a behavioural task based on high-level features (e.g. animacy categorisation) (Richter et al., 2024). This task could plausibly have impacted the observed prediction effects by increasing attention to task-relevant, high-level features and decreasing task-based attention to low level feature mismatches (Auksztulewicz et al., 2017; Smout et al., 2019). Given that the present study’s behavioural task placed minimal demands on attention to the semantic features of individual stimuli, it seems unlikely that task demands alone were responsible for driving the differential impact of low-and higher-level features observed. Notably, recent imaging work has also suggested that neural effects associated with contextual expectations emerge at different times in visual processing stages, depending on the degree of contextual exposure (Ferrari et al., 2022). Specifically, Ferrari et al. (2022) found that, when exposed to a predictable context, expectation effects emerged in areas associated with higher-order visual processing (e.g. inferior frontal gyrus) comparatively earlier than they appeared in earlier visual areas. In the present study, each exposure to semantic contexts (i.e. RSVP blocks) was very short (<40 seconds). In the context of Ferrari et al.’s (2022) findings, it is possible that only higher-level visual features modulated prediction effects because participants did not have sufficient exposure to predictive contexts for effects in areas encoding early visual features.

Importantly, these results should be interpreted with caution, as the present paradigm was not optimised for feature-level analyses. For example, individual stimuli can vary widely in terms of decoding accuracy and the stimuli included in high versus low dissimilarity comparison groups are, by nature, not identical (Grootswagers et al., 2022; Grootswagers et al., 2019). Dissimilarity measures are not independent at different featural levels, as stimuli with similar semantic content often share many lower-level features. The dissimilarity measures used here were correlated, which makes it challenging to clearly disambiguate between the role of different features. However, if this correlation was strong enough to fully mask critical differences, no differences would have been expected between the evaluated feature levels.

Additionally, these exploratory analyses investigated the role of feature-level dissimilarity in cases where stimuli were semantically unexpected but not semantically expected. Analysis was confined to unexpected stimuli as, in this design, dissimilarity measurements between individual stimuli and other members of the same semantic category often lacked sufficient variability to facilitate informative statistical testing. This approach might explain why higher-level features were implicated in decoding differences. Decoding differences between semantically unexpected and random stimuli emerged within the time window generally associated with the representation of higher-level image content such as semantic category information (e.g. animacy) (>200ms, Carlson et al., 2013). Given this timing, it is not necessarily surprising that subgroup-analyses indicated that decoding effects were more strongly linked to dissimilarity within the features represented at this time window. Notably, for semantically expected stimuli, decoding differences emerged much earlier (e.g. 125 ms) and continued throughout time windows associated with both low- and high-level feature encoding (Carlson et al., 2013). It is therefore plausible that the dissimilarity dynamics identified in the exploratory analyses may apply to semantic violation stimuli but may not fully account for the effects observed in semantically expected stimuli. This possibility could not be adequately explored in the present paradigm but remains an important topic for future work.

Our RSVP paradigm employed a semantic prediction manipulation where participants were briefly exposed to stimuli belonging to common, broad semantic categories. These stimuli were not optimised to facilitate investigation of the role of lower-level versus higher-level features, but future research could explore this question in detail by using stimulus sets with linearised low- and or high-level features. Previous studies have implemented predictive relationships using many different statistical relationships (Feuerriegel et al., 2021), and it is unclear whether the results identified here are generalisable to different operationalisations of semantic predictive relationships. It is also unclear the extent to which the learned predictive relationships were implicit or explicit, though participants often reported being aware that some blocks mainly presented stimuli from a single semantic category. Future work should aim to explore whether the results identified in the current study generalise to other types of predictive relationships, and the extent to which awareness may modulate observed prediction effects.

Overall, the results of this project provide novel insight into the neural dynamics associated with semantic-level predictive relationships. Different types of prediction violations were found to have distinct modulatory effects on neural responses. Exploratory analyses identified measures of semantic dissimilarity, rather that low-level visual feature dissimilarity, as key factors modulating the expression of neural prediction effects. Considered cumulatively, these results provide novel insight into the neural implementation of predictive processing.

## Funding

M.J.M. is supported by an Australian Research Council Discovery Early Career Research Award (ARC DE240100327). J.B.M. was supported by a National Health and Medical Research (NHMRC) Investigator Grant (Leadership 3; GNT2010141).

## Competing Interests

The authors report no competing interests

## Author Contribution Statement

In line with the CReDiT taxonomy, MJM was responsible for conceptualisation, formal analysis, methodology, software, supervision, visualisation, and writing – original draft. PD was responsible for formal analysis, methodology, software, visualisation, and writing – review & editing. XJO was responsible for data curation and prepared an early draft of a portion of this project for a student project (Writing – original draft). JBM was responsible for conceptualisation, supervision, and writing – review & editing.

## Supplemental Material

**Supplementary Table 1:** Summary of the main reported decoding analyses. ‘Training Data’ corresponds to the stimulus types used to train the classifier, whereas ‘Testing Data’ refer to the stimulus types used to test the model. Where testing data are not reported, models were trained and tested on the same stimuli. ‘N Epochs’ refers to the number of epochs (per participant) used to train or test each classifier. ‘Onset’ indicates the first time at which at least three sequential timepoints were significantly above chance. Onsets are not reported for analyses in which no timepoints survived corrections for multiple comparisons. ‘End’ refers to the last timepoint at which at least three sequential timepoints were above chance. In cases where ‘End’ is not reported, decoding accuracy remained above chance until the end of the relevant epoch. ‘Peak Accuracy’ is the maximum mean classifier accuracy, and ‘Peak Time’ is the time at which this occurred. ‘N BF > 3’ reports the number of timepoints at which decoding accuracy was above chance (BF > 3) between -100 and 1000ms from stimulus onset.

| Training Data | Testing Data | Figure | N Epochs | Onset | Peak Accuracy | Peak Time | N BF > 3 |
| --- | --- | --- | --- | --- | --- | --- | --- |
| *Semantic Expectancy Models:* |  |  |  |  |  |  |  |
| Random |  | 2 | 188 (80 - 216) | 82 | 56.55% | 195 | 197 |
| Random | Semantically Expected | 2 | 288 (60 - 485) | 66 | 55.57% | 125 | 250 |
| Random | Semantically Unexpected | 2 | 130 (44 - 199) | 78 | 57.00% | 203 | 239 |
| *Low-Frequency Violation Models:* |  |  |  |  |  |  |  |
| Random |  | 3 | 188 (79 - 216) | 78 | 56.80% | 195 | 187 |
| Random | Low-Frequency Violations | 3 | 130 (44 - 199) | 78 | 55.73% | 203 | 224 |

## Notes

https://osf.io/s6k7u/

### Competing Interest Statement

The authors have declared no competing interest.

